# Processes within the subspaces leading to changes in performance and keeping it unchanged

**DOI:** 10.64898/2026.03.23.713586

**Authors:** Sayan Deep De, Mark L. Latash

**Author notes:** Corresponding author: Sayan Deep De Department of Kinesiology Rec.Hall-268N, The Pennsylvania State University University Park, PA 16802, USA.

## Abstract

We explored processes within and orthogonal to the solution space (uncontrolled manifold, UCM) as superposition of fast random walk (RW) and slow drifts during multi-finger force production. Healthy participants performed two-hand (using the index and middle fingers per hand) accurate total force production task with different initial sharing of the force between the hand. After 5 s, visual feedback was manipulated – kept for both force and sharing, for only one of those variables, or turned off. The subjects tried to continue “doing what they have been doing” for 55 s. Trajectories both along and orthogonal to the UCM for total force showed fast RW and slow drifts. The diffusion plots confirmed persistent RW within the first 0.2 s and anti-persistent RW after 0.5 s. Persistent RW was similar across visual feedback conditions and larger orthogonal to the UCM. Its Hurst index correlated between the UCM and orthogonal to the UCM direction across participants. Anti-persistent RW depended strongly on visual feedback. Drift magnitude and characteristic time depended strongly on visual feedback, being similar along and orthogonal to the UCM. We conclude that RW destabilizes the state of the system thus encouraging exploration of nearby states over short time intervals and contributes to its stability over larger time intervals. Visual feedback plays a more important role in structuring stability of performance compared to the explicit task formulation. RW exploration promises new insights into the organization of stability in abundant systems and a potential biomarker for clinical studies.

**Highlights:**

- Stability during a multi-element action is structured in a feedback-specific way;
- Random walk and drift characteristics of force depend not on the task but on salient feedback;
- Random walk destabilizes steady state within a short range and stabilizes it within a long range;
- The control of an action encourages exploration but limits its range.

## Introduction

All natural human movements involve multiple contributing elements that can be explored at different levels of analysis. For example, reaching movements have been explored in the joint configuration spaces, prehensile movements in the spaces of force and moment vectors produced by the digits, whole-body movements in spaces of muscle activation, etc. (reviewed in Latash et al. 2007; Latash 2021). This fact has been framed as the problem of motor redundancy (Bernstein 1947) or the bliss of motor abundance (Gelfand and Latash 1998; Latash 2012). Within the latter approach, the abundance of elemental variables has been viewed as contributing to the dynamical stability of task-specific salient performance variables and explored, in particular, using the framework of the uncontrolled manifold (UCM) hypothesis (Scholz and Schöner 1999). While characteristics of data distributions within the solution space for a salient performance variable (the UCM) have shown sensitivity to neurological disorders and effects of practice (reviewed in Latash 2021), time processes within the UCM have not been explored until recently.

In a recent study of prolonged accurate production of the total force (F_TOT_) generated by a set of fingers, we described two processes that happen within the UCM: A random walk (RW) with the characteristic times of ≈ 50-100 ms, and drifts with the characteristic times of 5-15 s (De et al. 2025a). In that experiment, visual feedback on the F_TOT_ magnitude was always present, while the feedback on the coordinate along the UCM was shown only over the first 5 s and then turned off. The subjects were always instructed to “continue doing what you have been doing.” The timing characteristics of the RW suggested a major role of spinal circuitry, with possible contributions from both reflex feedback loops and intra-spinal loops such as recurrent inhibition (cf. Hultborn et al. 2004). The timing characteristics of the drift were similar to those reported for drifts in F_TOT_ in the absence of visual feedback on its magnitude (Vaillancourt and Russell 2002; Ambike et al. 2015; Cuadra et al. 2021).

In the current study, we explored the RW and drift characteristics along the UCM and orthogonal to the UCM space (ORT) during multi-finger total force (F_TOT_) production task using manipulations of visual feedback that could provide continuous feedback on the coordinate along one of these spaces (Z_UCM_ or Z_ORT_), both, or none. As in the previous study (De et al. 2025), the subjects had complete visual feedback on both Z_UCM_ and Z_ORT_ over the first 5 s of each trial, were asked to produce the same F_TOT_ magnitude (reflected in Z_ORT_) using different sharing of F_TOT_ between the index and middle finger pairs of both hands (reflected in Z_UCM_), and then continue “doing the same” over the next 55 s when the visual feedback was manipulated.

Figure 1 presents schematically our study using a two-effector F_TOT_ production task. The solution space (UCM) is shown with the slanted solid line, and the ORT space with the dashed line. We assume that an initial state of the system (the black circle) is associated with different stability properties along the UCM and ORT, shown schematically with two potential fields, shallow along the UCM (ϕ_UCM_) and steep along the ORT (ϕ_ORT_). This simple schematic leads to a set of specific hypotheses.

**Figure 1.**
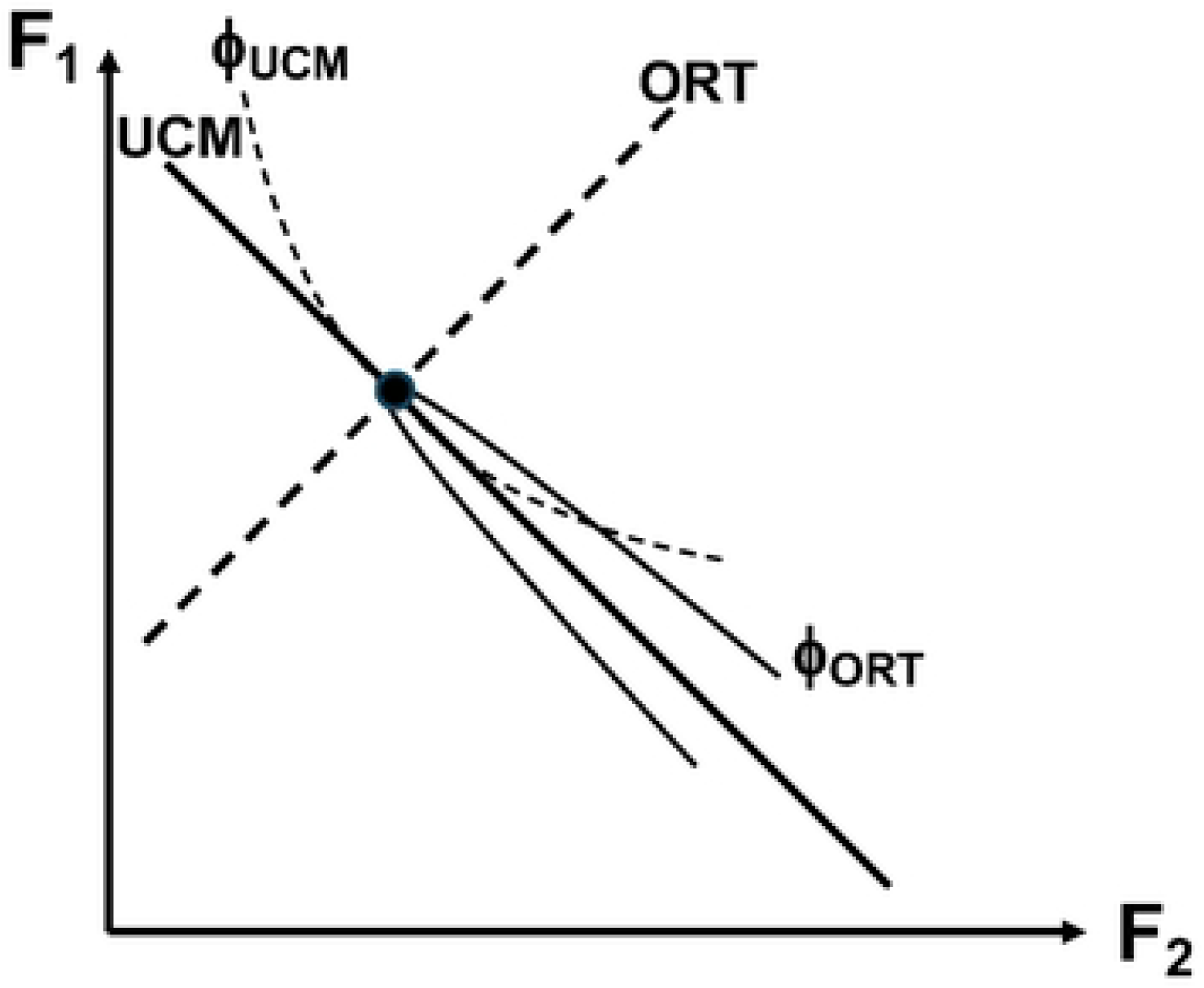
A schematic representation of a two-effector total force (F) production task. The uncontrolled manifold (UCM) is shown with the slanted solid line, and the ORT space with the dashed line. Different stability properties along the UCM and ORT for a state of the system along the UCM (the black circle) are shown schematically with two potential fields, shallow along the UCM (ϕUCM) and steep along the ORT (ϕ_ORT_).

Since the instructed performance variable was always the same (F_TOT_), we hypothesized that both RW and drift would be smaller along the ORT as compared to the UCM, reflecting the higher stability along the ORT (Hypothesis-1). RW has been invoked as a means of exploration of the solution space in search of globally preferred solutions during the processes of development and rehabilitation (Roth et al. 2023). No such hypotheses have been offered for the ORT, which makes experimental exploration of Hypothesis-1 necessary. We also expected processes along the ORT to be faster than along the UCM, also as a reflection of the different stability properties (Hypothesis-2; see Ambike et al. 2016; Reschechtko et al. 2015). Note that RW can correspond to purely random processes defining each step, be persistent or anti-persistent (Mandelbrot and van Ness 1968). In the latter case, the deviations from the initial coordinate are expected to accumulate more slowly, reflecting a degree of stability along that coordinate. Since the UCM is less stable (see also Martin et al. 2009, 2019), we expected the RW to show stronger anti-persistence along the ORT as reflected by the Hurst exponent in the diffusion plot analysis (Hypothesis-3; cf. Collins and DeLuca 1993). Earlier studies of drifts in the total force have suggested that, during steady-state tasks, these drifts (within the ORT, by definition) originated within the UCM and resulted from a weak coupling between the UCM and ORT (Ambike et al. 2016; Reschechtko and Latash 2017). Hence, we explored possible correlations between the main characteristics of the RW and drifts in the two spaces, in particular when no visual feedback was available.

## Methods

### Participants

Thirteen healthy adults participated in the experiment (7 males and 6 females; age: 23.7 ± 5.9 years, mean ± standard deviation). Their average mass was 75 ± 13.8 kg, and average height was 1.7 ± 0.1 m. All participants were right-handed according to self-reported everyday hand usage during writing and eating, had normal or corrected-to-normal vision, and reported no history of neurological or musculoskeletal disorders that could affect hand function. All participants gave written informed consent in accordance with procedures approved by the Office for Research Protections of the Pennsylvania State University.

### Equipment

Participants were seated in a comfortable chair with back support and positioned their forearms on wooden supports attached to a rigid table. Vertical finger forces were recorded using force sensors (Nano-17, ATI Industrial Automation, Garner, NC, USA). The sensors were mounted within rigid frames that allowed comfortable placement of the fingers while maintaining a consistent hand posture throughout the trials. Participants produced force using the index and middle fingers of both hands. The contact surfaces of the sensors were covered with sandpaper to prevent slipping. Signals from the sensors were amplified and sampled at 500 Hz using a 16-bit data acquisition board (National Instruments, Austin, TX, USA).

Visual feedback was displayed on a 19-inch monitor positioned approximately 0.6 m in front of the participant at eye level. Custom software developed in LabVIEW controlled data acquisition, and feedback display. The experimental setup is schematically showed in Fig. 2A.

**Figure 2.**
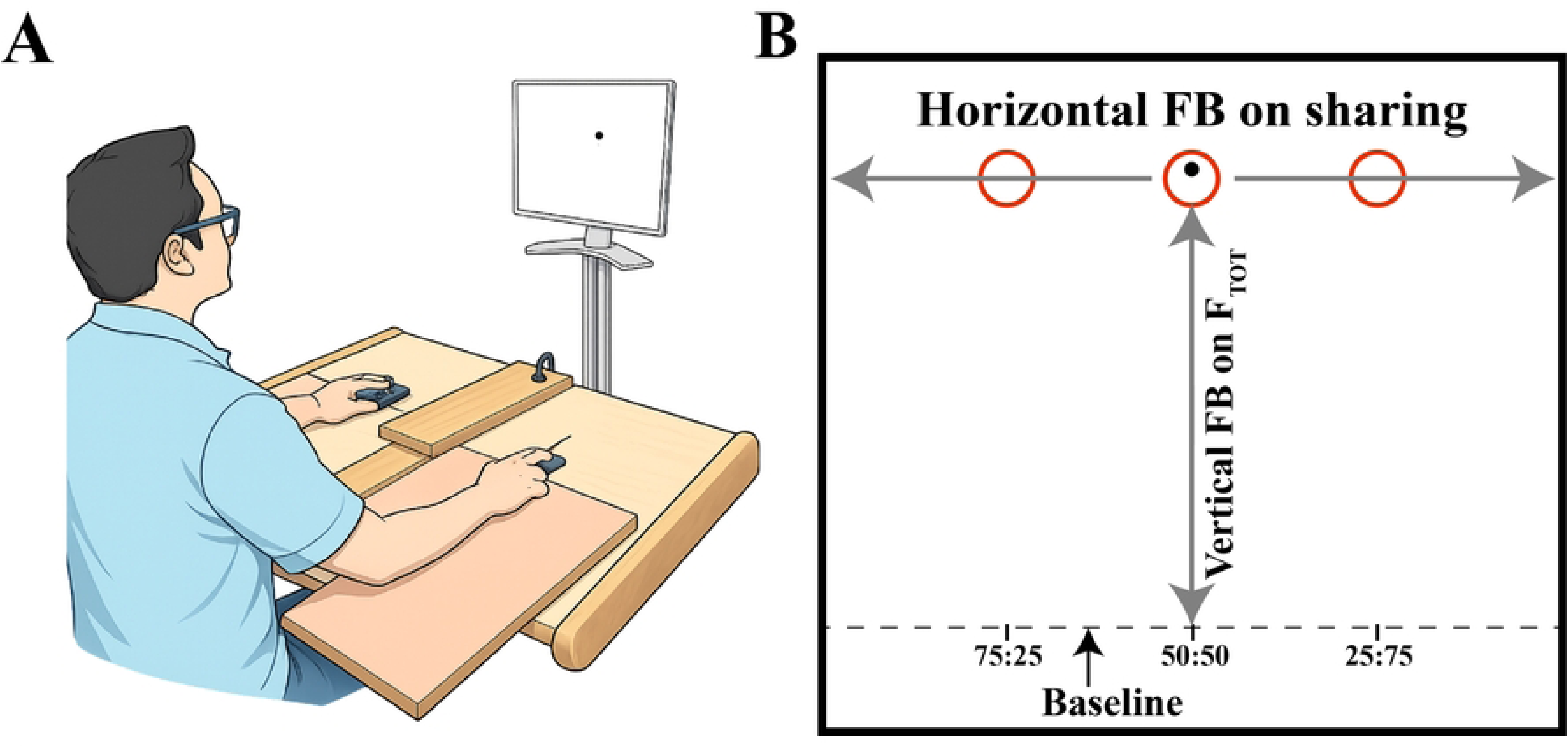
A: Schematic illustration of the experimental setup. Participants produced vertical forces with the index and middle fingers of both hands using force sensors mounted in rigid slots. B: Organization of the feedback display. The vertical axis represented the total force magnitude (F_TOT_), whereas the horizontal axis represented the sharing index (SI), defined as the percentage of force produced by the right hand. Targets corresponding to the three sharing configurations (25:75, 50:50, and 75:25) are illustrated as red circles.

### Experimental Procedure

#### Maximum voluntary contraction (MVC)

At the beginning of the session, participants performed three maximum voluntary contraction (MVC) trials. During these trials, participants gradually increased their pressing force following an auditory “go” signal and attempted to reach their maximal force within approximately 3 s while receiving visual feedback of the total force produced by all four fingers. The highest force achieved across trials was defined as the MVC and was used to scale the force targets in the subsequent experimental trials.

#### Experimental task

Participants performed a 60-s accurate force production task while producing a target total force corresponding to 15% MVC. The visual display represented two task variables simultaneously: the total force magnitude (F_TOT_) and the sharing of force between the two hands (sharing index, SI). The vertical axis of the display corresponded to the total force magnitude, whereas the horizontal axis represented the relative contribution of the two hands. Targets were displayed corresponding to three SI patterns: 25:75, 50:50, and 75:25. SI corresponds to the percentage of force produced by the right hand. The feedback screen is illustrated in Fig. 2B.

Each trial began with a short period (5 s) during which full visual feedback was available, allowing participants to position the cursor within the target region. Following this initial phase, specific components of visual feedback were removed depending on the experimental condition. Four feedback conditions were tested:

- FB_F_ – feedback on F_TOT_ only;
- FB_S_ – feedback on SI only;
- FB_B_ – feedback on both F_TOT_ and SI; and
- FB_N_ – no visual feedback.

Participants were instructed to maintain the cursor within the target and continue producing force in the same manner (“continue doing what you have been doing”) throughout the 60-s trial. One trial was performed for each combination of sharing configuration and feedback condition. There were 30-s rest intervals between trials to avoid fatigue. None of the participants reported fatigue.

### Data Processing

All analyses were performed offline using custom scripts in MATLAB and R. Raw force signals were visually inspected for artifacts and low-pass filtered using a fourth-order zero-lag Butterworth filter with a cutoff frequency of 50 Hz. Forces produced by the fingers of each hand were summed to obtain left-hand (F_L_) and right-hand (F_R_) forces. The total force was computed as: F_TOT_ = F_L_ + F_R_.

To remove edge effects and transient adjustments at the beginning and end of the trial, analyses were restricted to the interval between 7 s and 59 s.

To illustrate the temporal structure of the performance, Fig. 3 shows representative time profiles of the signals along the UCM (Z_UCM_) and ORT (Z_ORT_) directions for a typical trial under the 25:75 sharing condition and the FB_N_ condition. In this example, both coordinates show slow drifts superimposed on faster fluctuations. The definitions of Z_UCM_ and Z_ORT_ metrics are presented in detail in the next section. To visualize the fast component independently of the slow drift, we computed the lagged-difference signals using a 1-s lag:

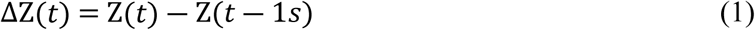

**Figure 3.**
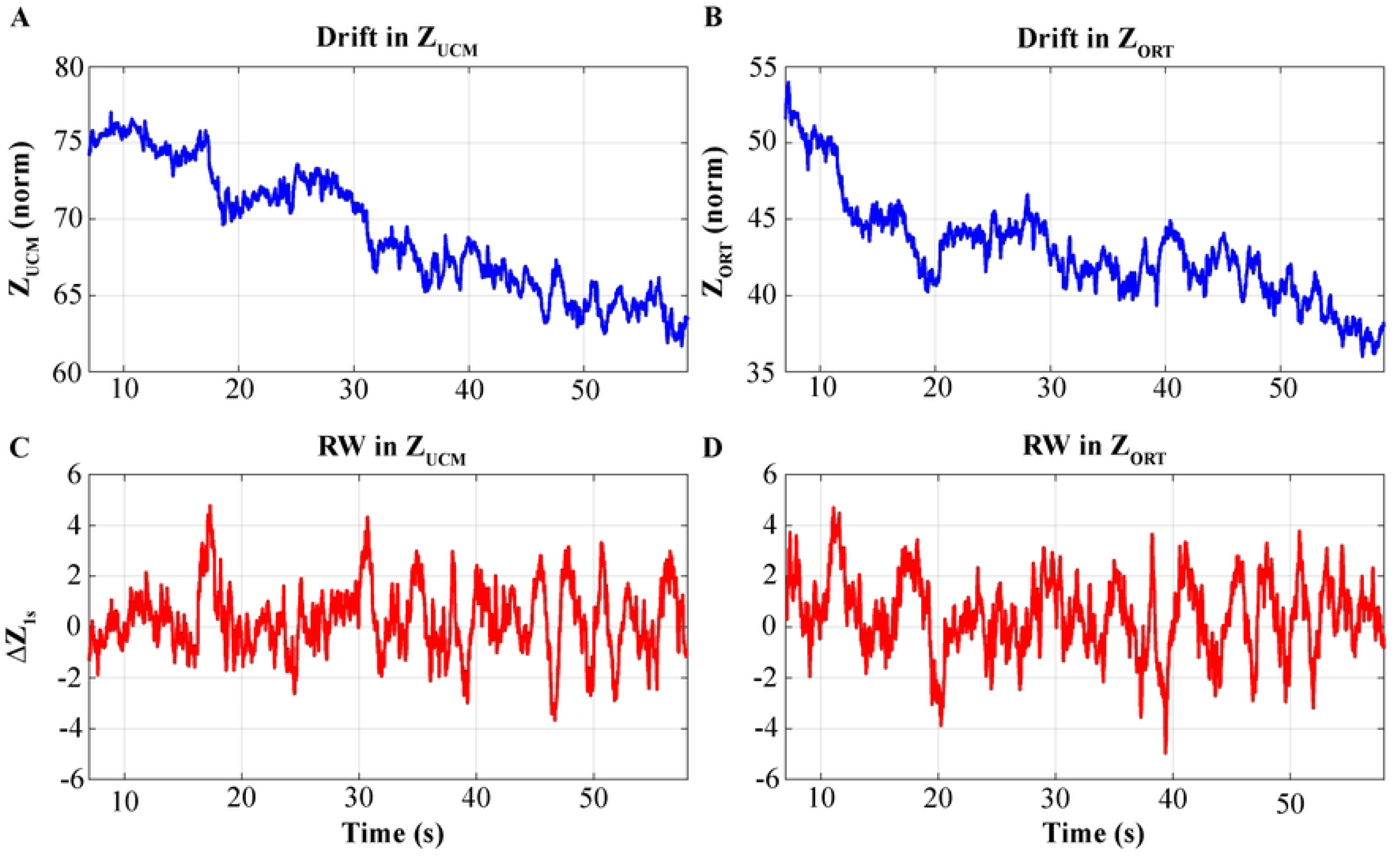
Representative time profiles of the task coordinates during a typical trial performed under the 25:75 sharing and no-feedback (FB_N_) condition. A: Time profile of Z_UCM_ illustrating slow drift superimposed on faster fluctuations. B: Time profile of Z_ORT_ showing a similar combination of slow drift and fast fluctuations. C: Lagged-difference signal computed from ZUCM using a 1-s lag (ΔZ_UCM_). D: Lagged-difference signal computed from Z_ORT_ using a 1-s lag (Δ_ZORT_). The lagged-difference transformation suppresses slow drift while preserving the faster fluctuations associated with the random-walk component.

where Z denotes either Z_UCM_ or Z_ORT_. Th transformation (1) suppresses slow trends while preserving the high-frequency fluctuations associated with the RW component. The lagged-difference method has been described in detail in our earlier study (De et al., 2025a). The original trajectories and their corresponding 1-s lagged-difference signals are illustrated in Fig. 3 for the UCM (left panels) and ORT (right panels) for F_TOT_.

#### Task coordinates

To separate processes affecting F_TOT_ from those preserving it, the behavior was analyzed using two task-related variables derived from the forces produced by the left and right hands. To make values along the ORT (Z_ORT_) and along the UCM (Z_UCM_) commensurable, we used the following metrics. In the initial condition, for the 50:50 sharing, the distance along the UCM to each of the force axes was assumed to be 50 units (corresponding to the percentage of the sharing index). The distance from the origin of coordinates along the ORT direction to the initial state for the 50:50 sharing was also assumed to be 50 units to preserve geometric similitude (Z_ORT_ = 50, Figure 4). These metrics were kept across conditions. Since the task required the production of F_TOT_ = 15% of MVC, this normalization made 15%MVC equivalent to 50 normalized force units.

**Figure 4.**
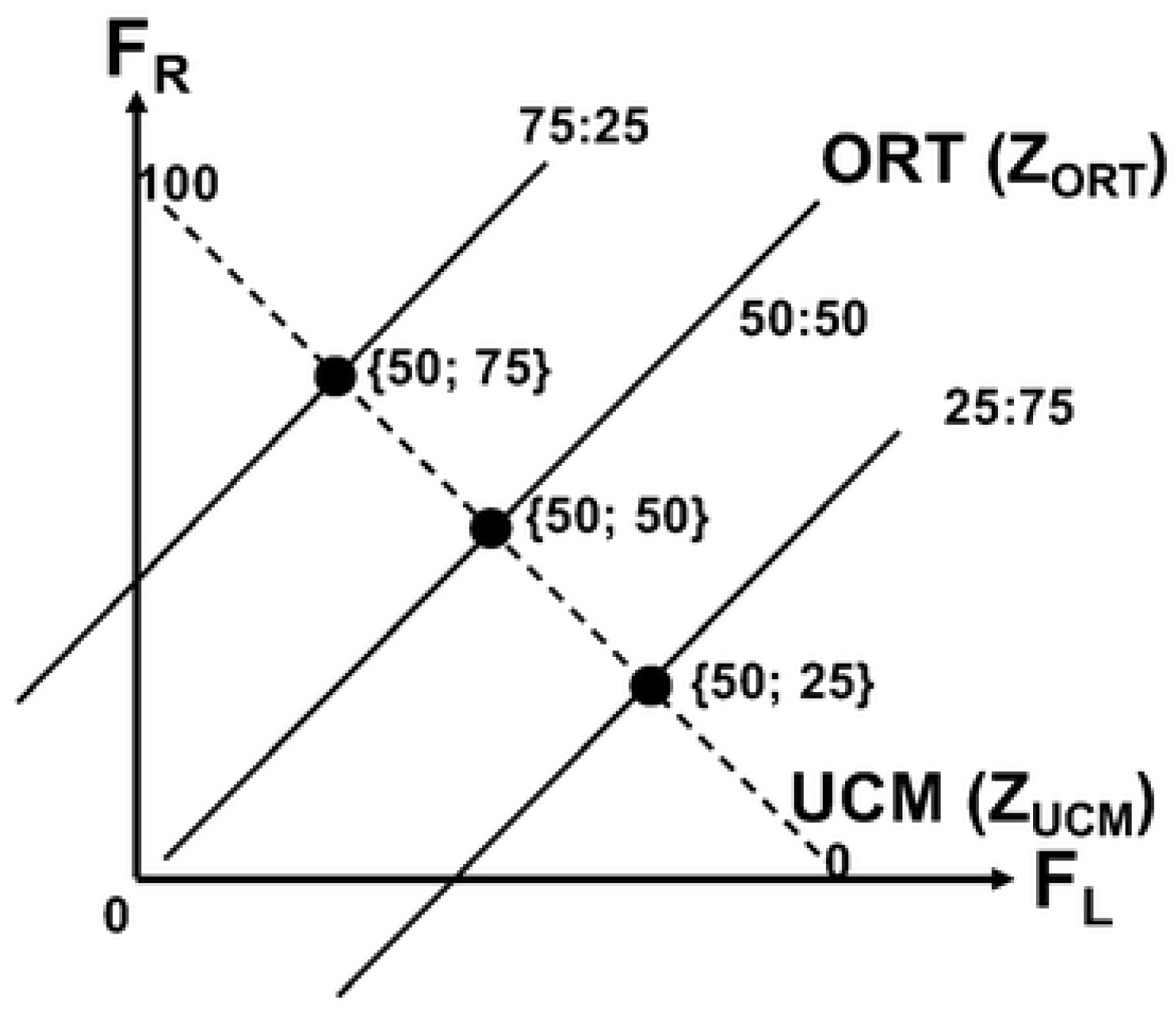
Schematic illustration of the three initial states corresponding to different coordinates along the uncontrolled manifold (UCM, dashed line) and orthogonal (ORT, solid lines) directions. The total force requirement was always the same assumed to be 50 normalized units; the three sharing pattern requirements (75:25, 50:50, and 25:75 shown on top of the ORT lines) corresponded to coordinates 75, 50, and 25 along the UCM resulting in three points with the coordinates {50; 75}, {50; 50}, and {50; 25}. Note that normalization of force to 50 units makes coordinates along the UCM and ORT commensurable.

Z_UCM_ = 0 corresponds to F_TOT_ produced entirely by the left hand, whereas Z_UCM_ = 100 corresponds to F_TOT_ produced entirely by the right hand. Z_UCM_ = 50 represents equal sharing between the two hands. Note that changes in the task variable, F_TOT_ magnitude, are reflected in the deviations along the ORT space (Z_ORT_ coordinate) while changes in the sharing without a change in F_TOT_ are reflected in the deviations along the UCM (Z_UCM_ coordinate). Note also that Z_ORT_ was always measured along the ORT direction across the sharing patterns.

#### Drift analysis

Slow changes in the trajectories along each coordinate will be referred to as drift processes. Drift indices were computed separately for Z_ORT_ and Z_UCM_ using the following indices:

- Peak-to-peak magnitude (PP) – defined as the difference between the maximum and minimum values of the coordinate within the analysis window.
- Trial drift (TRIAL) – defined as the difference between the initial and final coordinate values within the trial.
- Drift time constant (τ_50_) – defined as the minimal time interval required for the coordinate to change by 50% of the PP magnitude.

To reduce the influence of fast fluctuations, trajectories were smoothed using a moving average window (5 s; implemented as a sample-based window using the median sampling interval) prior to estimation of τ_50_. Note that this procedure was used in the drift analysis only, and raw trajectories were used for the random walk analysis (see later).

#### Histogram drift analysis

To characterize how behavior evolved over the course of a trial, we constructed histograms of the relevant task variable within the analysis window (7–59 s). For the UCM dimension, this was done using the sharing index (SI), with values assigned to bins spanning 0–100% SI so that the resulting distributions reflected the amount of time spent at different sharing values. Similar histograms were also computed for the ORT direction, where the distributions reflected the amount of time spent at different force levels. Histograms were computed separately for each feedback condition and sharing configuration, and bin heights were normalized by the number of participants.

For illustrative purposes, only the FB_F_ and FB_S_ conditions are shown in Figure 5. Figure 5A illustrates the drifts along the UCM dimension under the FB_F_ condition, whereas Fig. 5B shows the drifts along the ORT direction under the FB_S_ condition. Note the predominance of drifts toward 50:50 in conditions with uneven sharing (25:75 and 75:25) in Fig. 5A. Note also predominance of F_TOT_ drift toward lower magnitudes across the sharing patterns in Fig. 5B.

**Figure 5.**
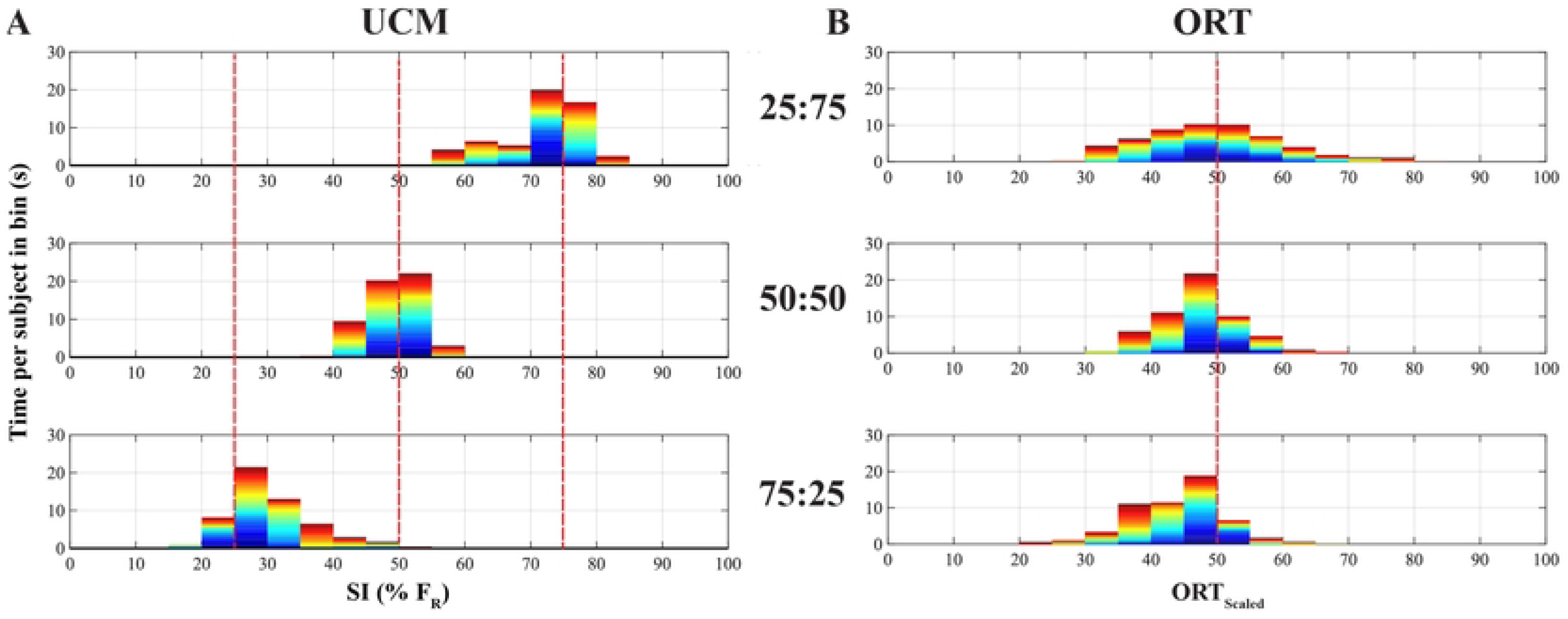
Histograms of the distributions of the task coordinates across subjects for the three sharing conditions, 25:75, 50:50, and 75:25. A: Distribution of Z_UCM_ values illustrating drift along the UCM direction during the condition with feedback on total force only (FB_F_). B: Distribution of Z_ORT_ values illustrating drift along the ORT direction during the condition with feedback on sharing only (FB_S_). Dark blue colors correspond to early time intervals, and brighter (red) colors correspond to times closer to the end of the trial. The vertical red lines show the initial coordinates for each task. Note the different drift directions in A and similar drift directions in B for different sharing conditions.

### Random walk analysis

Fast fluctuations superimposed on the drift component were analyzed as a random-walk process using diffusion analysis. For Z_ORT_ coordinate and Z_UCM_ separately, the mean squared displacement was computed as a function of the time increment (Δt):

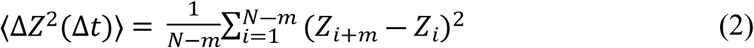

where Z denotes either Z_ORT_ or Z_UCM_, *N* is the number of samples in the analyzed interval, and *m* is the lag corresponding to Δt = *m* · dt (dt = sampling interval). The resulting diffusion plots were represented in log–log coordinates. Linear regressions were fit to two regions of the diffusion plot:

- Short window: 0–0.2 s; and
- Long window: 0.5–1.5 s.

These windows were selected based on the previous study (De et al. 2025a) to reflect the two regimes consistently seen across subjects when the slopes of the diffusion plot changed. The slopes of these regressions (α) were used to compute the Hurst exponent (H = α/2), which characterizes the temporal structure of the fluctuations:

H = 0.5 indicates classical Brownian motion,

H > 0.5 indicates persistent behavior,

H < 0.5 indicates anti-persistent behavior.

#### RW frequency and power estimation

Across conditions and sharing configurations, the distributions of cycle durations derived from zero crossings of the velocity signal were visually similar, showing no systematic differences in their overall shape. Therefore, for a compact summary, the distributions were averaged across feedback conditions and sharing ratios, and group-level frequency statistics were computed from the pooled data.

Oscillatory fluctuations in the analyzed frequency range were evaluated for the UCM and ORT spaces by estimating signal band-power from the corresponding time series.

#### Statistics

Data are presented as means ± standard deviations unless stated otherwise. Prior to parametric analyses, the distributions of all variables were examined for normality using Shapiro–Wilk tests and visual inspection of residuals. When appropriate, variables were log-transformed to improve normality, and rank transformations were applied when distributions remained non-normal. Sphericity of repeated-measures factors was evaluated using Mauchly’s test, and when violated the Greenhouse–Geisser correction was applied.

Drift characteristics along the two task coordinates (Z_ORT_ and Z_UCM_) were quantified using three indices: peak-to-peak magnitude (PP), cumulative drift over the trial (TRIAL), and the time required to reach 50% of the total drift (τ_50_). These metrics were analyzed using repeated-measures ANOVA with Feedback Condition (four levels: FB_F_, FB_S_, FB_B_, FB_N_) and Sharing configuration (three levels: 25:75, 50:50, 75:25) as within-subject factors. When needed, PP and τ_50_ variables were analyzed after log transformation, while cumulative drift measures were rank-transformed to better satisfy distributional assumptions. Significant ANOVA effects were explored using Tukey-adjusted pairwise comparisons.

Random-walk (RW) characteristics were evaluated using scaling exponents (H) derived from the diffusion analysis. Two time windows were examined separately: a short window (0–0.2 s) and a long window (0.5–1.5 s). Because H values did not always satisfy parametric assumptions, factorial effects of Feedback and Sharing were tested using Aligned Rank Transform (ART) ANOVA, a non-parametric procedure that enables analysis of multifactor repeated-measures designs while preserving the ability to evaluate interactions. Short- and long-window scaling values were also compared to the theoretical value of 0.5 using Wilcoxon signed-rank tests.

The power values were analyzed using an ART ANOVA with factors Space (SI, ORT) and Sharing (25:75, 50:50, 75:25). Holm-corrected pairwise comparisons were used to examine differences between spaces and sharing ratios when significant effects were detected.

To examine relationships between the RW processes in the two task subspaces, Pearson’s correlation analysis was performed between the Hurst exponents computed for the Z_ORT_ and Z_UCM_ coordinates.

The level of statistical significance was set at p < 0.05. All statistical analyses were performed using the R programming language within the RStudio environment.

## Results

Across conditions, participants successfully performed the 60-s accurate force-production task and, after the initial period with full visual feedback, their behavior systematically depended on which feedback components remained available. Specifically, there were slow changes (drifts) and faster, random-walk–like, fluctuations in the coordinates along both the uncontrolled manifold (Z_UCM_) and its orthogonal complement (Z_ORT_). Drifts of Z_UCM_ were most prominent when visual feedback on sharing was absent (FB_F_ and FB_N_), whereas drifts of Z_ORT_ were most evident when feedback on F_TOT_ was absent (FB_S_ and FB_N_). When both feedback signals were continuously available (FB_B_), trajectories in both coordinates remained close to their initial values with no consistent drift. In the sections below, we describe these effects separately for the drift and random-walk (RW) components within each subspace, beginning with qualitative illustrations and then presenting the statistical analyses.

### Processes affecting the force magnitude (F_TOT_)

#### Drifts that change F_TOT_

Under the feedback conditions FB_S_ and FB_N_ leading to large drifts in the coordinate along the ORT space (Z_ORT_), F_TOT_ showed predominantly drifts to lower force magnitudes. These are illustrated as histograms for the FB_S_ condition in Figure 5B. This Figure summarizes the data across subjects. Note that the drifts were similar across the two uneven sharing conditions (25:75 and 75:25) while they were smaller for the 50:50 sharing.

Drift in the coordinate along the ORT space (Z_ORT_) was quantified using peak-to-peak magnitude (Z_ORT_-PP; log-transformed), cumulative drift over the trial duration (Z_ORT_-TRIAL; rank-transformed), and the time required to reach 50% of the drift (Z_ORT_-τ50; log-transformed). All three metrics depended primarily on the feedback condition and showed only modest dependence on the prescribed initial sharing ratio, with no evidence for a sharing-by-condition interaction.

For Z_ORT_-PP (Fig. 6A), ANOVA revealed significant main effects of Condition (F_(2.44, 29.30)_ = 102.84, p < 0.001, η²₍_G_₎ = 0.65) and Sharing (F_(1.68, 20.12)_ = 6.93, p = 0.007, η²₍_G_₎ = 0.06), while the Sharing × Condition interaction was not significant (p = 0.174). Post-hoc comparisons showed that drift magnitude differed across sharing ratios mainly due to the reduced drift in the 50:50 sharing relative to 75:25 (Tukey-adjusted p = 0.0029). Across feedback conditions, Z_ORT_-PP was significantly larger in FB_S_ or FB_N_ conditions (feedback on sharing or no feedback at all) than in FB_F_ and FB_B_ conditions (p < 0.0001), indicating that force feedback either alone or in combination with sharing feedback attenuated drift along Z_ORT_.

**Figure 6.**
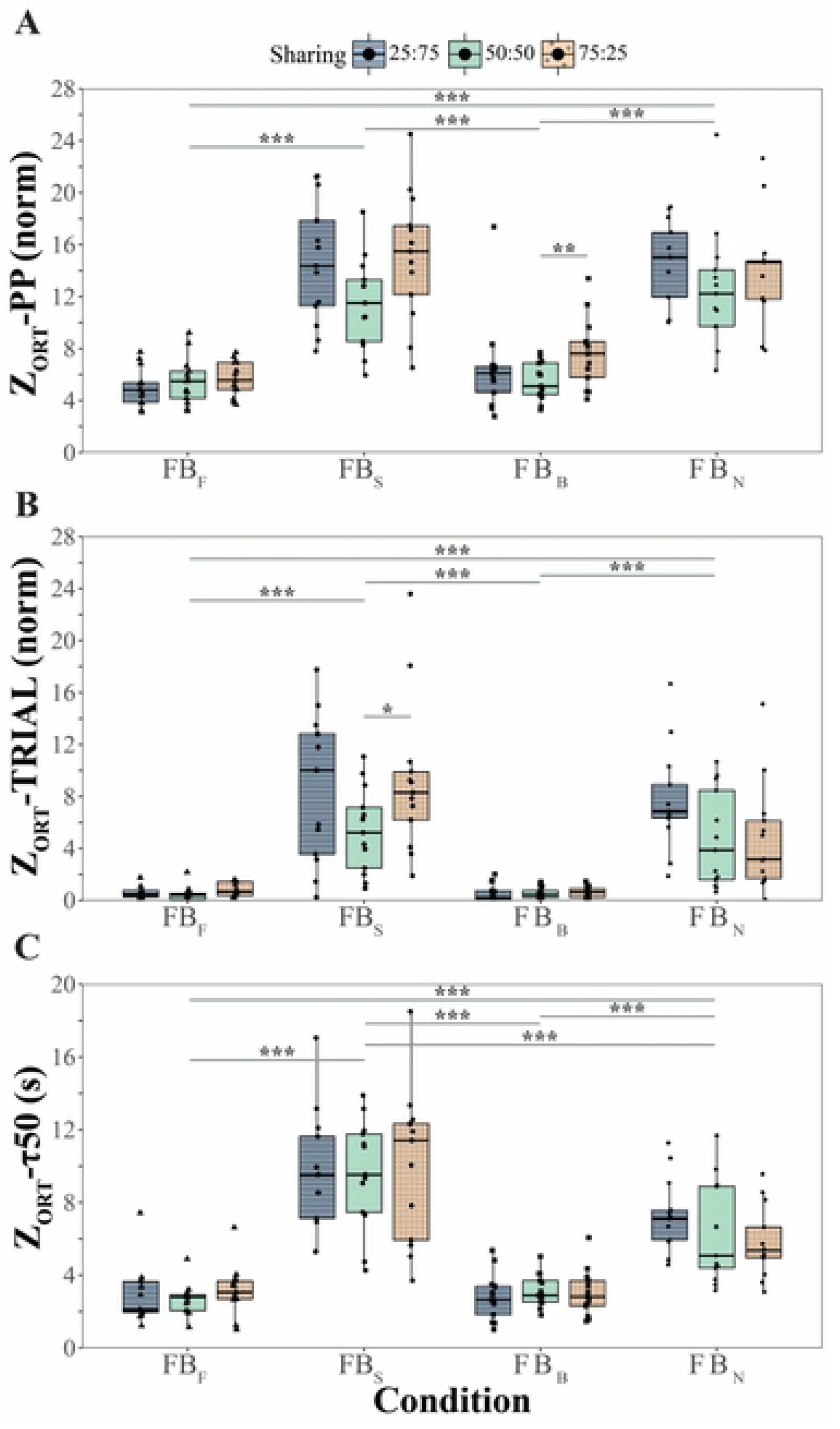
Quantification of drifts along the ORT direction (Z_ORT_). A: Peak-to-peak magnitude of the drift (Z_ORT_-PP). B: Total change over the trial duration (Z_ORT_-TRIAL). C: Time required to reach 50% of the peak-to-peak magnitude (τ50). Box-and-whisker plots show individual participant data for the three sharing patterns and four feedback conditions. Note the relatively large drift magnitudes along the ORT direction across the experimental conditions where force feedback was turned off (FB_S_ and FB_N_).

A very similar pattern emerged for cumulative drift (Z_ORT_-TRIAL; Fig. 6B). The main effect of Condition was robust (F_(2.17, 26.00)_ = 72.77, p < 0.001, η²₍_G_₎ = 0.66), while the effect of Sharing showed only a trend (p = 0.057), and the Sharing × Condition interaction was not significant (p = 0.177). Z_ORT_-TRIAL was significantly larger under feedback on sharing FB_S_ or no feedback at all, FB_N_, conditions than FB_F_ and FB_B_ conditions (p < 0.0001).

The timing of the drift, captured by Z_ORT_-τ50 (Fig. 6C), was also dominated by Feedback condition (F_(2.16, 25.87)_ = 73.45, p < 0.001, η²₍_G_₎ = 0.65). Neither the main effect of Sharing (p = 0.973) nor the Sharing × Condition interaction (p = 0.441) reached significance. Z_ORT_-τ50 was the longest under FB_S_, shorter under FB_N_ (p < 0.005), and shortest when F_TOT_ feedback was provided either alone (FB_F_) or in combination with sharing feedback (FB_B_).

#### Random walk affecting F_TOT_

The pooled distributions of the cycle duration for the UCM and ORT coordinates are illustrated in Fig. 7. Overall, there was no significant difference between the rapid fluctuations along both coordinates, which occurred within a similar frequency range. When all trials were combined, the median oscillation frequency of Z_UCM_ (reflecting the sharing index, SI) was 19.23 Hz (quartiles: 10.87–35.71 Hz). A comparable distribution was observed for Z_ORT_ (reflecting F_TOT_ changes), with a median frequency of 15.63 Hz (quartiles: 10.00–31.25 Hz).

**Figure 7.**
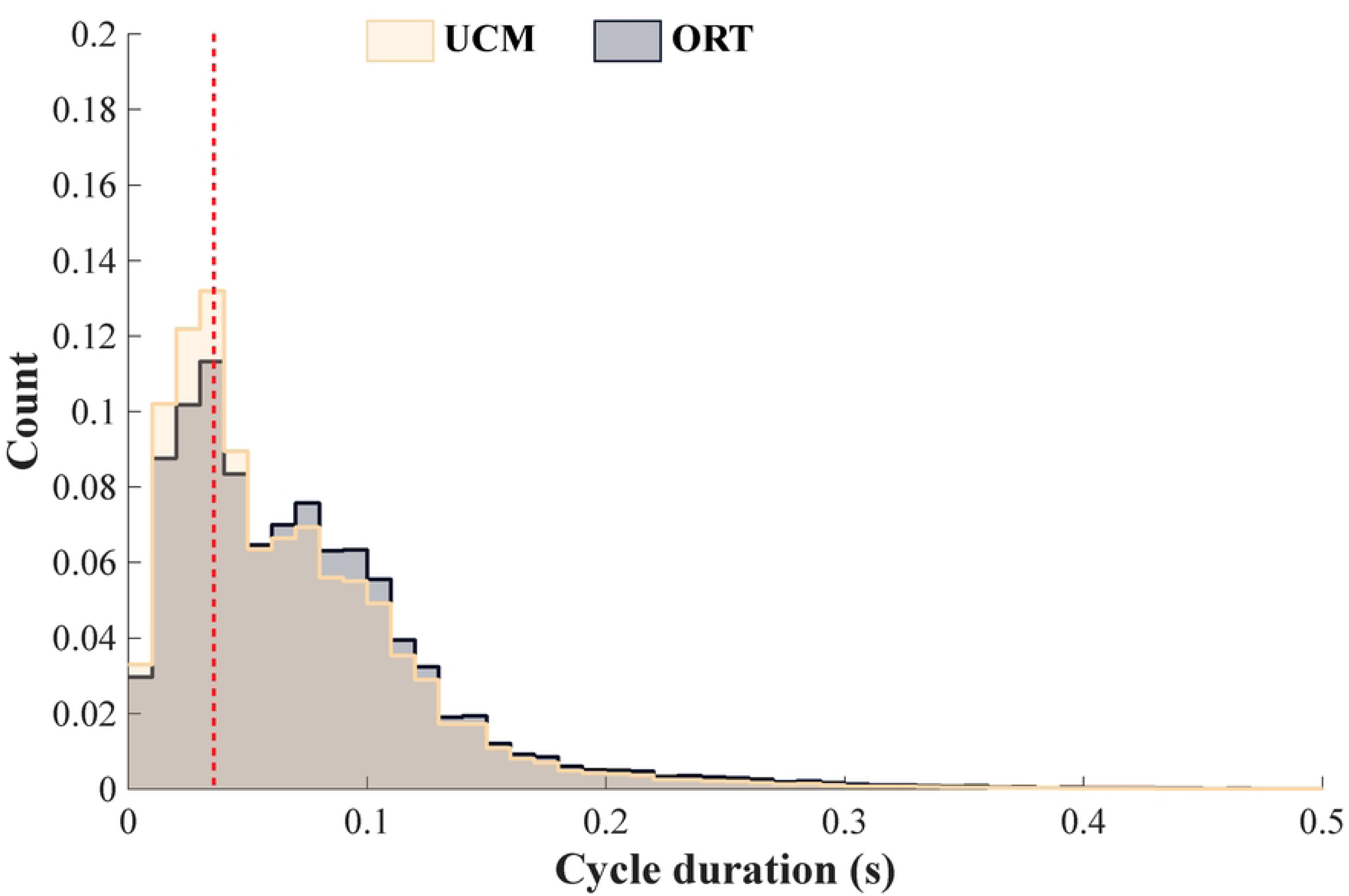
Pooled histograms of cycle duration for the UCM and ORT coordinates computed from zero crossings of the corresponding velocity signals. Cycle durations were grouped into bins of width 0.01 s. The distributions are pooled across all subjects, feedback conditions, and sharing configurations. The y-axis shows relative frequency (normalized count), computed as the proportion of all detected cycles falling within each bin. The dashed vertical line indicates the time point with the maximum count; note that this value was the same for both UCM and ORT. Note the similar distributions for the two coordinates, with most values concentrated below 0.1 s.

RW explored for Z_ORT_ revealed a clear separation between the two time scales. In the short window (0-0.2 s), the Hurst exponent (H_Short_) was consistently greater than 0.5 (mean ± SD: 0.67 ± 0.07), indicating persistent dynamics. These results are illustrated in Figure 8. A Wilcoxon signed-rank test confirmed that H_Short_ was significantly larger than 0.5 (V = 91, p = 0.0008). In contrast, within the long window, 0.5-1.5 s, H_Long_ showed values well below 0.5 (0.26 ± 0.04), consistent with anti-persistent behavior. A Wilcoxon test confirmed that H_Long_ was significantly smaller than 0.5 (V = 0, p = 0.0008). Thus, RW along the ORT exhibited persistence over short time scales and anti-persistence over long time scales.

**Figure 8.**
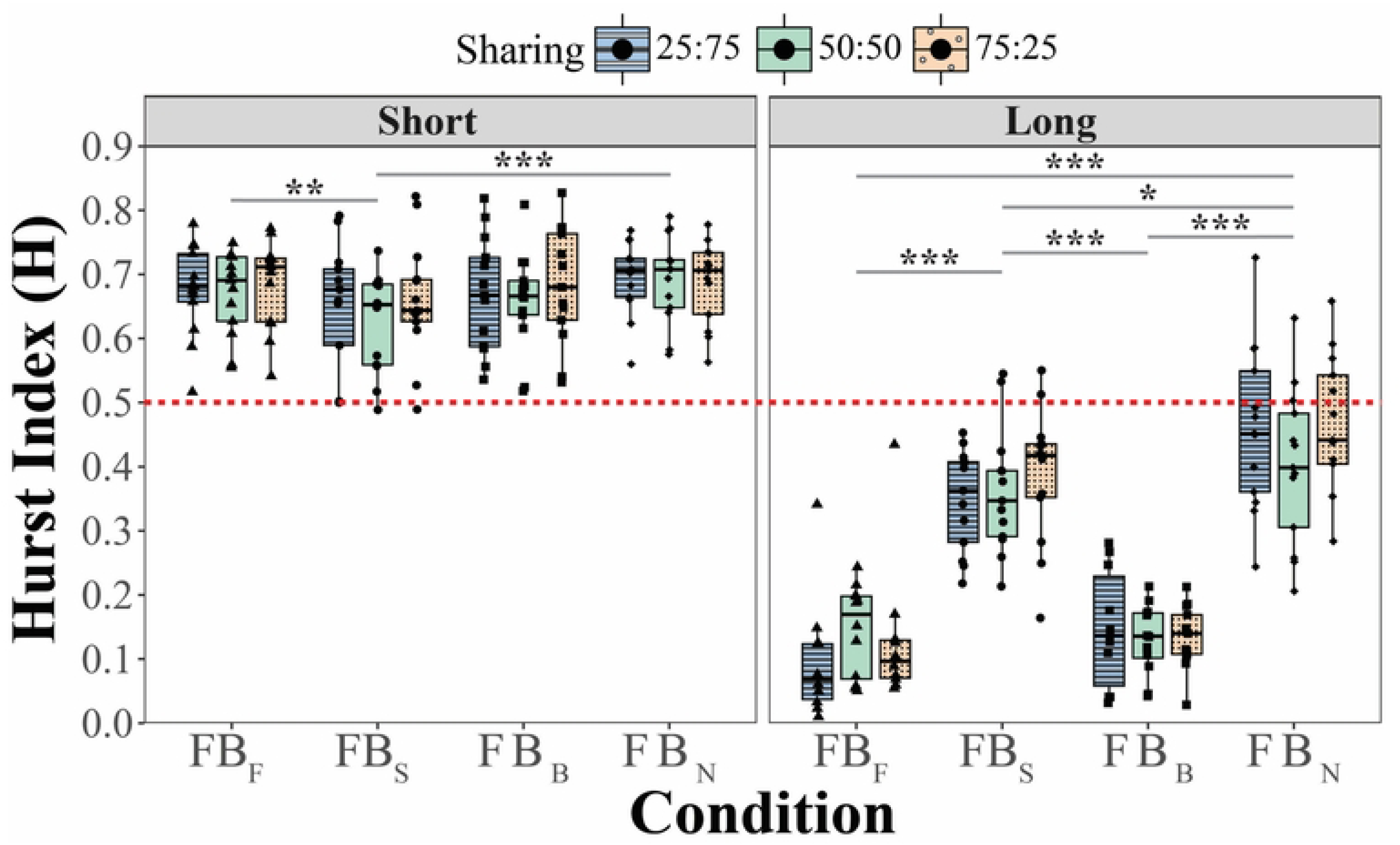
Random-walk characteristics along the ORT direction. Short-window Hurst exponent (H_Short_) computed within the 0–0.2 s interval and long-window Hurst exponent (H_Long_) computed within the 0.5–1.5 s interval. Box-and-whisker plots show individual participant values for the different sharing configurations and feedback conditions. Note the persistent dynamics at short time scales similar across the feedback conditions (H_Short_ > 0.5) and anti-persistent dynamics (H_Long_ > 0.5) at longer time scales different across the feedback conditions.

Given the clear quantitative separation between the short- and long-window scaling exponents, the effects of feedback condition and sharing were examined separately for H_Short_ and H_Long_. H_Short_ values showed only modest variation across task conditions. An ART ANOVA with Condition and Sharing as factors revealed a significant main effect of Condition (F_(3,132)_ = 8.61, p < 0.001; η²₍_P_₎ = 0.16), whereas the effect of Sharing and the interaction were not significant. Pairwise comparisons indicated that there were smaller H_Short_ values under FB_S_ than under FB_F_ (p = 0.005) and FB_N_ (p < 0.001), while the remaining comparisons were not significant.

A markedly stronger effect emerged for H_Long_. As seen in the right panel of Figure 8, H_Long_ values were largest when no feedback was available (FB_N_), somewhat smaller under sharing feedback (FB_S_), and substantially smaller whenever force feedback was present (FB_F_ and FB_B_). An ART ANOVA confirmed a strong main effect of Condition (F_(3,132)_ = 120.91, p < 0.001; η²₍_P_₎ = 0.73), whereas the effect of Sharing and the interaction were not significant. Post-hoc comparisons showed that both force-feedback conditions (FB_F_ and FB_B_) differed strongly from the conditions lacking force feedback (FB_S_ and FB_N_) (all p < 0.001), while the two force-feedback conditions (FB_F_ and FB_B_) did not differ from each other. Likewise, the two conditions without force feedback produced significantly larger H_Long_ values.

### Processes that did not change F_TOT_

#### Drifts along the UCM

Under the feedback conditions FB_F_ and FB_N_ leading to large drifts in the coordinate along the UCM space (Z_UCM_), the sharing index showed predominantly drifts toward the 50:50 pattern. There were, however, exceptions. The effects of drifts are illustrated as histograms for the FB_F_ condition in Figure 5A. This Figure summarizes the data across subjects. Note that the drifts were larger under the two uneven sharing conditions (25:75 and 75:25) as compared to the 50:50 sharing.

Drift along the uncontrolled manifold (Z_UCM_) was quantified using the same three indices: peak-to-peak drift magnitude (Z_UCM_-PP), cumulative drift over the trial (Z_UCM_-TRIAL), and the time required to reach 50% of the peak-to-peak drift (Z_UCM_-τ50). Across all metrics, drift along the UCM was strongly modulated by the availability of visual feedback on sharing (see Figure 9).

**Figure 9.**
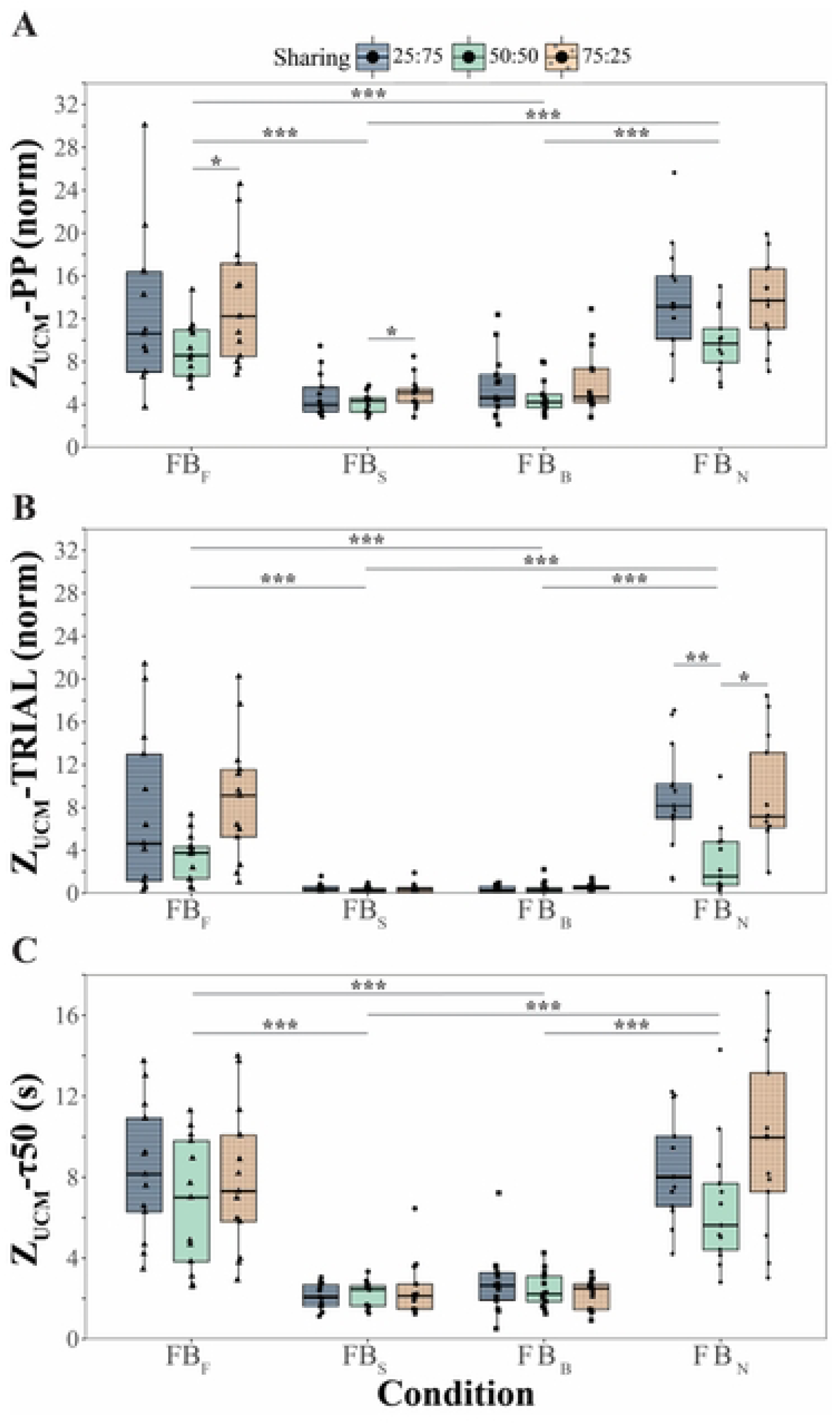
Quantification of drift along the uncontrolled manifold (Z_UCM_). A: Peak-to-peak magnitude of the drift (Z_UCM_-PP). B: Total drift over the trial duration (Z_UCM_-TRIAL). C: Time required to reach 50% of the peak-to-peak magnitude (Z_UCM_-τ50). Box-and-whisker plots show individual participant data across sharing configurations and feedback conditions. Note the relatively large drift magnitudes along the ORT direction across the experimental conditions where sharing feedback was turned off (FB_F_ and FB_N_).

Both Z_UCM_-PP (log-transformed; Fig. 9A) and Z_UCM_-TRIAL (rank-transformed; Fig. 9B) showed robust main effects of Condition (Z_UCM_-PP: F_(2.40, 28.75)_ = 86.68, *p* < 0.001, η²₍_G_₎ = 0.56; Z_UCM_-TRIAL: F_(2.41, 28.92)_ = 86.46, *p* < 0.001, η²₍_G_₎ = 0.65), indicating substantial differences in drift magnitude across feedback conditions. In both measures, the drift was markedly larger when visual feedback on sharing was absent (FB_F_, FB_N_) than when sharing feedback was available (FB_S_, FB_B_). No significant Sharing × Condition interactions were observed (all *p* > 0.07).

In addition, both Z_UCM_-PP and Z_UCM_-TRIAL exhibited significant main effects of the initial sharing ratio (Z_UCM_-PP: F_(1.63, 19.61)_ = 5.32, *p* = 0.019, η²₍_G_₎ = 0.08; Z_UCM_-TRIAL: F_(1.66, 19.95)_ = 7.32, *p* = 0.006, η²₍_G_₎ = 0.1). Across conditions, drift magnitudes were generally smallest for the symmetric 50:50 sharing and larger for the asymmetric sharing configurations.

The temporal drift characteristic, Z_UCM_-τ50 (log-transformed, Fig. 9C) demonstrated a strong main effect of Condition (F_(1.92, 22.98)_ = 85.10, *p* < 0.001, η²₍_G_₎ = 0.66), with larger values in conditions lacking sharing feedback, when the other drift metrics were also higher. In contrast, the main effect of the initial sharing ratio did not reach significance (*p* = 0.105), and no interactions were detected, indicating that the time scale of the drift along the UCM was primarily determined by the feedback condition rather than by the initial sharing configuration.

#### Random walk along the UCM

Random-walk along the UCM showed a clear separation between the short and long time scales (Figure 10). In the short time window, the Hurst exponent was consistently greater than 0.5 (mean ± SD: 0.66 ± 0.07), indicating persistent behavior of the RW. A Wilcoxon signed-rank test confirmed that H_Short_ was significantly larger than 0.5 (V = 91, p = 0.00083). In contrast, the long time window yielded values well below 0.5 (0.26 ± 0.03), consistent with anti-persistent dynamics. A Wilcoxon test confirmed that H_Long_ was significantly smaller than 0.5 (V = 0, p = 0.00083). A direct comparison between the two regimes showed that H_Short_ was significantly larger than H_Long_ (V = 91, p = 0.0017; mean difference = 0.40), indicating a pronounced change in scaling across the time scales.

**Figure 10.**
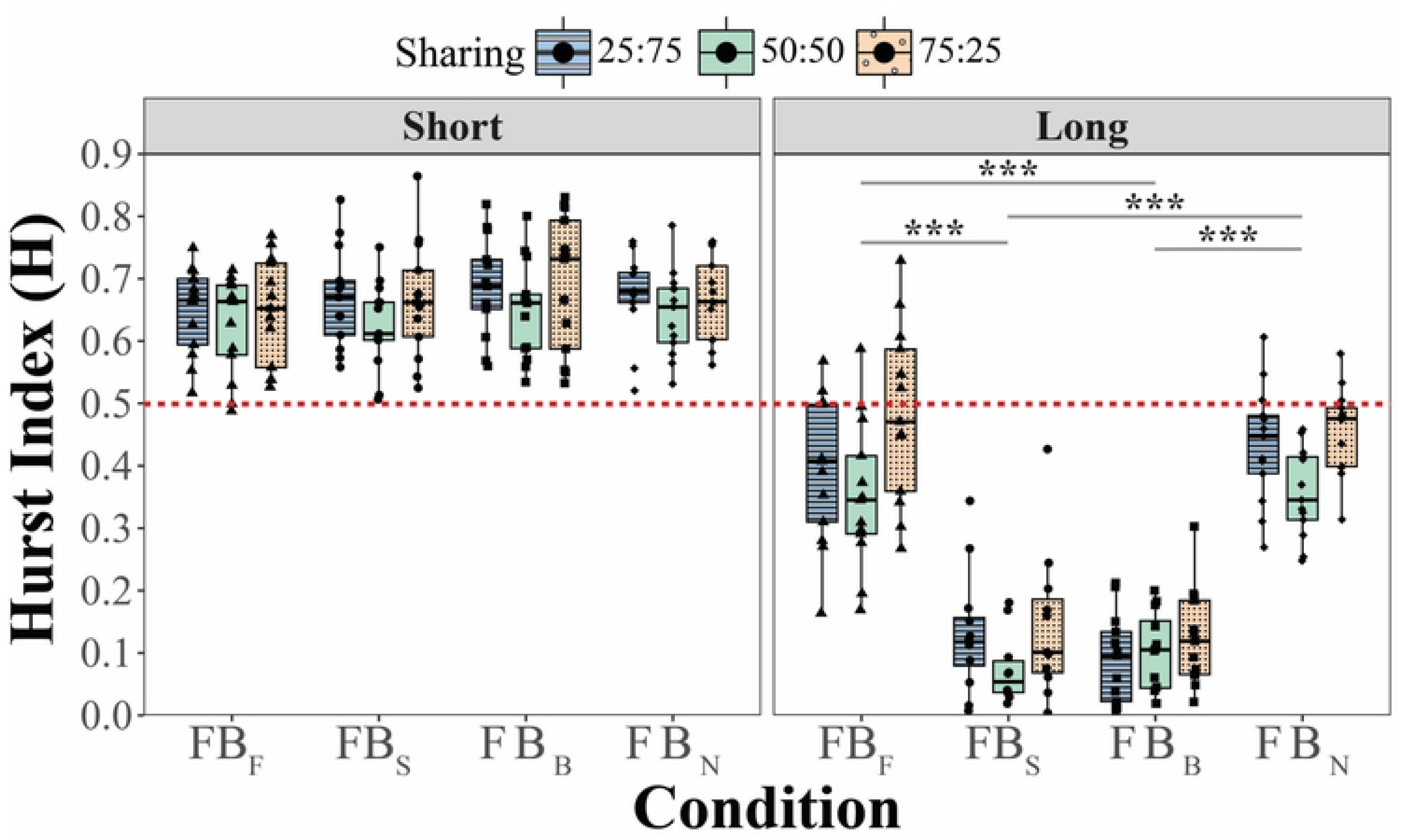
Random-walk characteristics of fluctuations along the UCM direction. Short-window Hurst exponent (H_Short_) estimated for the 0–0.2 s interval and long-window Hurst exponent (H_Long_) estimated for the 0.5–1.5 s interval. Note the similar pattern of persistent short-term dynamics across the feedback conditions (H_Short_ > 0.5) and anti-persistent long-term dynamics (H_Long_ < 0.5) depending on the available feedback.

Given this clear quantitative separation, the effects of task factors were examined separately for H_Short_ and H_Long_. H_Short_ showed moderate differences across conditions and sharing configurations. An ART ANOVA revealed significant main effects of Condition (F_(3,132)_ = 3.76, p = 0.013, η²₍_P_₎ = 0.08) and Sharing (F_(2,132)_ = 9.01, p < 0.001, η²₍_P_₎ = 0.12), while their interaction was not significant. Pairwise comparisons indicated that the condition with feedback on both force and sharing (FB_B_) produced larger H values than the condition with force feedback only (p = 0.013). In addition, the symmetric sharing configuration (50:50) yielded smaller H values than both asymmetric configurations (25:75 and 75:25; p ≤ 0.001).

A stronger and more systematic pattern emerged for the long time window data. The right panel of Figure 10 shows that H_Long_ values were largest when sharing feedback was absent (FB_F_ and FB_N_) and markedly smaller when sharing feedback was available (FB_S_ and FB_B_). An ART ANOVA confirmed significant main effects of both Condition (F_(3,132)_ = 121.97, p < 0.001, η²₍_P_₎ = 0.73) and Sharing (F_(2,132)_ = 9.86, p < 0.001, η²₍_P_₎ = 0.13), with no significant interaction. Post-hoc comparisons showed that the conditions without sharing feedback produced substantially larger H values than those with sharing feedback (all p < 0.001), while the two conditions within each pair did not differ from each other. In addition, the 50:50 configuration yielded lower H values than the asymmetric sharing ratios, consistent with the pattern observed in the short window.

Comparison of the H-values between the two spaces, UCM and ORT, showed significantly higher values along the ORT for H_Short_ (V = 80; p = 0.007) without significant differences in H_Long_ (p = 0.527). In other words, the persistent (destabilizing) effects over the short range were stronger along the ORT direction.

### Power estimation of fast fluctuations along the UCM and ORT

The band-power of the signal within the frequency range over 10 Hz differed across the UCM and ORT spaces and sharing ratios, although the differences were relatively small (see Figure 11). Overall, these differences depended on the sharing ratio, with the symmetric 50:50 condition associated with reduced Z_ORT_ band-power. The aligned rank transform ANOVA revealed significant main effects of Space (F_(_₁,₆₀_)_ = 18.67, p < 0.001) and Sharing (F_(_₂,₆₀_)_ = 3.85, p = 0.027), as well as a significant Space × Sharing interaction (F_(_₂,₆₀_)_ = 6.58, p = 0.003).

**Figure 11.**
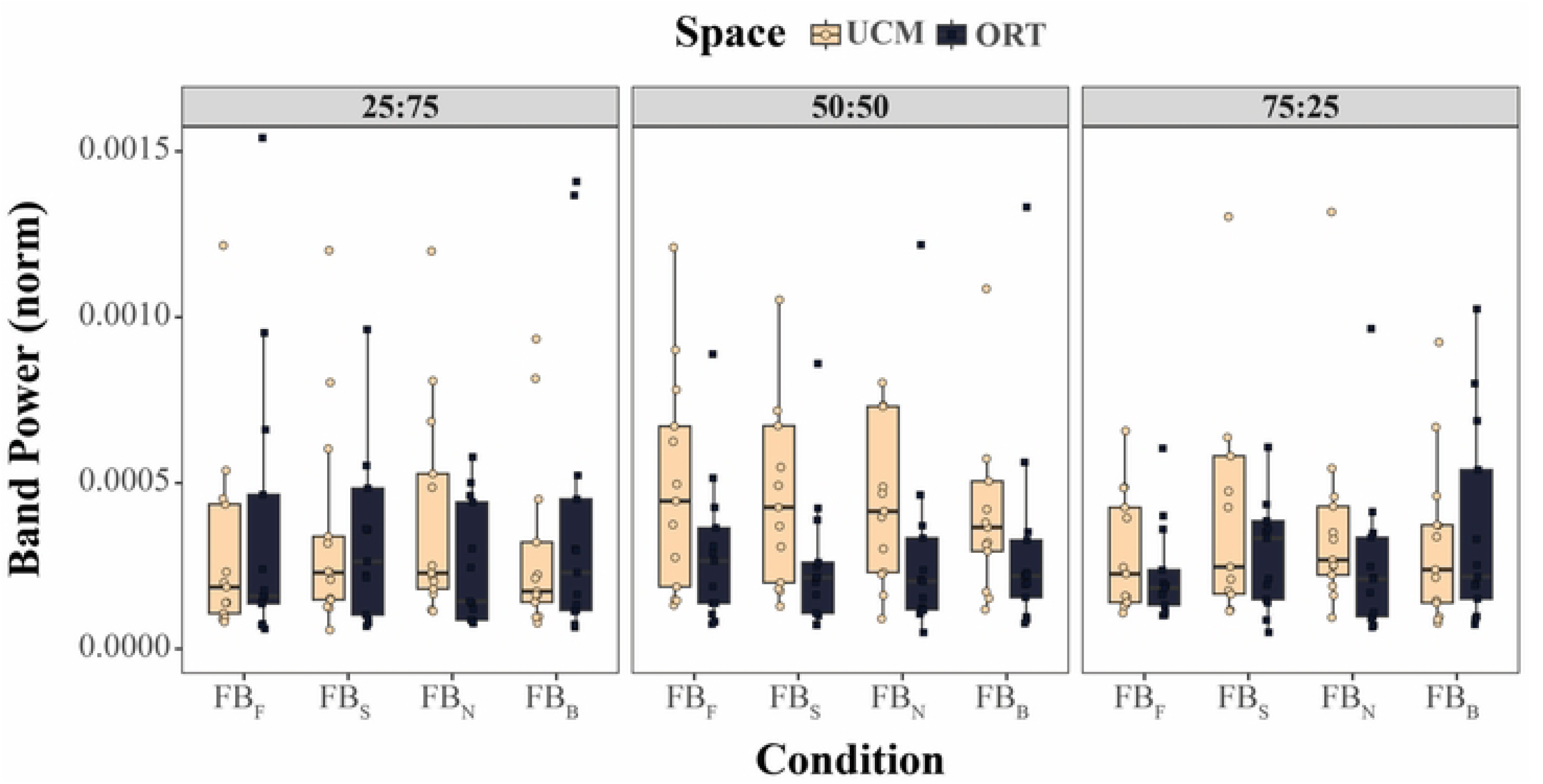
Band-power of fluctuations with the frequency >10 Hz for the UCM and ORT coordinates. Box-and-whisker plots show individual participant data for the three sharing patterns.

Holm-corrected pairwise comparisons showed that Z_ORT_ band-power exceeded Z_UCM_ band-power at 25:75 (p = 0.004), whereas Z_UCM_ tended to exceed Z_ORT_ at 50:50 (p = 0.051); no difference between spaces was observed at 75:25 (p = 0.113). Across sharing ratios, Z_UCM_ band-power did not differ significantly (all p > 0.05). In contrast, Z_ORT_ band-power was lower at 50:50 than at both 25:75 (p = 0.027) and 75:25 (p = 0.011).

### Comparison of RW between the UCM and ORT

Across conditions, the short-timescale Hurst exponents estimated along the ORT and UCM directions showed a very strong positive association across the participants. Pearson correlations were high for all conditions (FB_F_: r = 0.883; FB_S_: r = 0.883; FB_B_: r = 0.880; FB_N_: r = 0.868), indicating that participants who exhibited larger H_Short_ for Z_ORT_ also showed correspondingly larger H_Short_ for Z_UCM_. These results are illustrated in Figure 12, which shows the individual data and regression lines with equations. No such correlations were observed for H_Long_.

**Figure 12.**
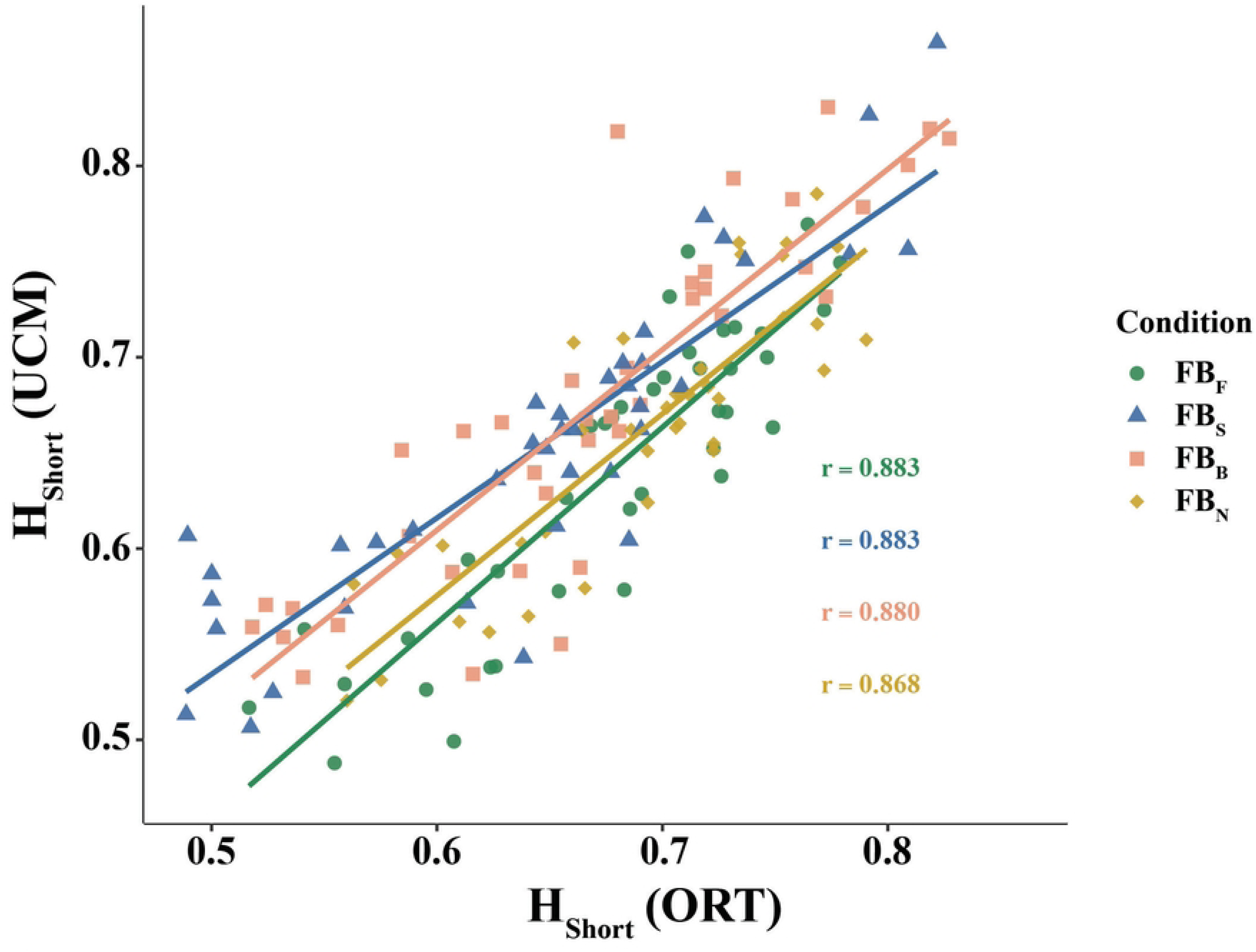
Short-time Hurst exponents (H_Short_) estimated along the ORT and UCM directions. The data for all subjects are shown separately for the four feedback conditions. Linear regression lines and the correlation coefficients are shown. Note the strong positive relation between the H_Short_ values along the ORT and UCM directions.

## Discussion

One of the main results of the study has been the demonstration that stability properties along the ORT and UCM spaces, as reflected by characteristics of both drifts and RW, are defined primarily by the available feedback, not by the explicit task formulation. The participants were always explicitly instructed to produce the same constant total force, F_TOT_, magnitude while the initial sharing pattern could vary. So, we hypothesized that, across all conditions, the UCM would be characterized by lower stability as compared to the ORT as reflected in the magnitude (Hypothesis-1, larger along the UCM) and speed (Hypothesis-2, larger along the ORT) of the drift and Hurst indices computed for the RW component (Hypothesis-3, smaller along the ORT). All three hypotheses have been confirmed when visual feedback on the force magnitude (Z_ORT_) was provided. However, they were all falsified when visual feedback on the sharing index (Z_UCM_) was provided.

A novel, potentially important, result is that the Hurst index was consistently over 0.5 corresponding to persistent RW for relatively small time windows of analysis and consistently under 0.5 corresponding to anti-persistent RW for larger windows of analysis. We observed a similar pattern in the previous study, but it was marginally significant (De et al. 2025). In the current set of experiments, this result was robust and observed across conditions and analyses. This result suggests that the RW component plays two roles. It encourages exploration of the solution space (cf. Roth et al. 2023) within a relatively small range while acting against major deviations from a preferred solution resulting in a non-trivial pattern of stability along both UCM and ORT discussed in more detail later.

Further, we discuss possible origins of the most unexpected findings, in particular a possibility that the presence or absence of visual feedback could lead to task reformulation by the neural controller. We also consider whether these results are specific to force production tasks in isometric conditions or could be expected across different types of tasks and external conditions.

### Factors that define the UCM

Since its introduction, the UCM hypothesis has been used as a tool to quantify stability of performance variables produced by large sets of elements (Scholz and Schöner 1999; reviewed in Latash et al. 2007; Vaz et al. 2019; Latash 2021). From the very first studies, the application of this toolbox demonstrated stabilization of different performance variables some of which were explicitly related to the task formulation while others were not. In particular, during accurate multi-finger force production tasks during pressing in isometric conditions, the participants demonstrated preferential stabilization of the moment of force in pronation-supination along the longitudinal axis of the hand/forearm, although this variable was not mentioned in the task formulation and the visual feedback was used to reflect the total force, not the total moment (Latash et al. 2001; Scholz et al. 2002).

By itself, the concept of UCM is theoretical as the solution space for the production of specific values or time profiles of a potentially salient performance variable. Its neural mechanisms have been linked to both feed-forward covaried changes in the elemental variables, likely based on practice (Goodman and Latash 2006) and feedback loops both within the central nervous system and from peripheral sensory endings (Latash et al. 2005; Martin et al. 2009, 2019). Several recent studies quantified the relative contributions of those mechanisms and showed the persistence of patterns of covariation seen early in the trial (likely reflecting feed-forward contributions) over tens of seconds (Abolins et al. 2025; De et al. 2025b). On the other hand, an important role of sensory feedback for stabilization of various performance variables has been confirmed in a number of studies. In particular, removing visual feedback on a performance variable leads to drifts in that variable in spite of the presence of natural somatosensory feedback (Vaillancourt and Russell 2002; Ambike et al. 2015). During isometric force and moment production tasks, the drift could reach over 30% of the initial magnitude of the performance variable, and the participants were unaware of those large errors in performance (Parsa et al. 2016, 2017). During tasks involving movement of an extremity or of the whole body, drifts of steady states are not readily seen, possibly due to the high resolution of position-related somatosensory signals. Such drifts, however, can be observed if the initial position is perturbed by an external device or by a self-generated action (Zhou et al. 2015; Rasouli et al. 2017).

One of the earlier studies used accurate cyclical total force (F_TOT_) production by a set of fingers with the help of the metronome and visual feedback on the F_TOT_ magnitude, which was turned off after the initial few seconds (Ambike et al. 2016). The study suggested the existence of two types of drifts with different characteristic times. Following the removal of the visual feedback, the midpoint of the cycle drifted to lower F_TOT_ magnitudes with the typical times of about 10 s, while the peak-to-peak force changes increased over much shorter times, ≈ 1 s.

Comparably fast drifts were confirmed in other studies with quick changes in the salient performance variable produced either by an external device or voluntarily (Wilhelm et al. 2013; Zhou et al. 2014). Those two types of drifts were hypothesized to originate from processes in spaces with different stability properties, the UCM (less stable, slow) and ORT (more stable, fast).

In our study, the explicit task formulation always emphasized the accurate production of a certain magnitude of F_TOT_ starting from different initial sharing patterns (SI), both reflected in the visual feedback. Based on the aforementioned hypothesis, we expected slow drifts along the UCM for F_TOT_ and fast drifts along the ORT. This hypothesis has been falsified. Indeed, drifts were pronounced along directions without visual feedback, but their speed (characteristic time, τ50) was comparable along the UCM and ORT. One possible interpretation of this observation is that the participant’s CNS reformulated the task depending on the available visual information and, as a result, the UCM and ORT directions switched when feedback for SI was available as compared to those when F_TOT_ feedback was available. An alternative interpretation is that the subjects associated different SI values with different total moment of force acting on the body in the frontal plane. This is feasible given the preferential stabilization of the total moment in both single-hand multi-finger tasks and two-hand tasks (Li et al. 2001; Scholz et al. 2002).

### Random walk and its potential roles

Random walk (RW) was originally developed as a mathematical abstraction of a one-dimensional process with equal probabilities of making a step with a standard magnitude from the current coordinate in one of the two possible directions. Further, this concept has been developed for multi-dimensional spaces and for processes with probabilities of stepping in a particular direction dependent on the previous step (Mandelbrot and van Ness 1968). If chances of stepping in the same direction are higher, the process is addressed as persistent; if they are lower, the process is addressed as anti-persistent. Using the concept of stability, persistent processes reflect unstable conditions and anti-persistent processes reflect different degrees of stability of the variable. These processes have been quantified using the so-called Hurst exponent (H) computed based on diffusion plots reflecting the deviation of the process from an initial state with time, with H > 0.5 corresponding to persistent (unstable) processes, and H < 0.5 – to anti-persistent (stable) processes.

The RW concept has been used in human movement studies across a variety of tasks and variables, from whole-body movements to eye movements (Collins and DeLuca 1993; Hausdorff et al. 1995; Kitazawa 2002; Shwetlick et al. 2025) including studies of the effects of aging and neurological disorders (Collins et al. 1995; Mitchell et al. 1995). It has been invoked as a potential exploratory mechanism important, in particular, during motor learning and motor rehabilitation (Roth et al. 2023).

In our experiment, across the visual feedback conditions and for processes along both the UCM and ORT, we consistently observed H > 0.5 over relatively short time intervals (H_Short_) and H < 0.5 over larger time intervals (H_Long_) when effects of corrections based on visual feedback could be expected. There were no effects of visual feedback conditions on H_Short_ and the values of H_Short_ were larger along the ORT compared to UCM. These observations suggest that H_Short_ reflects a basic process that is independent of the available visual feedback. Note that H_Short_ was estimated over a short time range (< 0.2 s) reflecting only a few steps of the RW process with the estimated average step duration of 0.03-0.07 s. Over this time interval, the deviation from the initial state increased in a persistent way thus contributing to the exploration of the space (cf.

Roth et al. 2023; De et al. 2025a). The larger values of H_Short_ along the ORT across the visual feedback conditions show that the instruction to the subject makes the destabilizing processes along the ORT direction stronger, an unexpected result given that many earlier studies emphasized higher stability along the ORT direction (reviewed in Latash 2021, 2024).

In contrast, H_Long_ showed significant effects of both visual feedback and direction of analysis (UCM vs. ORT). The small values of H_Long_ correspond to anti-persistent processes, i.e., those typical of movement in a stabilizing potential field. The different values and features of H_Short_ and H_Long_ suggest a non-monotonic potential field along both UCM and ORT with the slope of the anti-persistent (stabilizing) segment of the field larger along the direction with visual feedback. This is illustrated in Fig. 13 for a two-element F_TOT_ production task (cf. Fig. 1 in the Introduction). Note the short-range convex portion of the potential field amplifying deviations from the initial state (the black dot) and the long-range concave portion of the field acting against large deviations.

**Figure 13.**
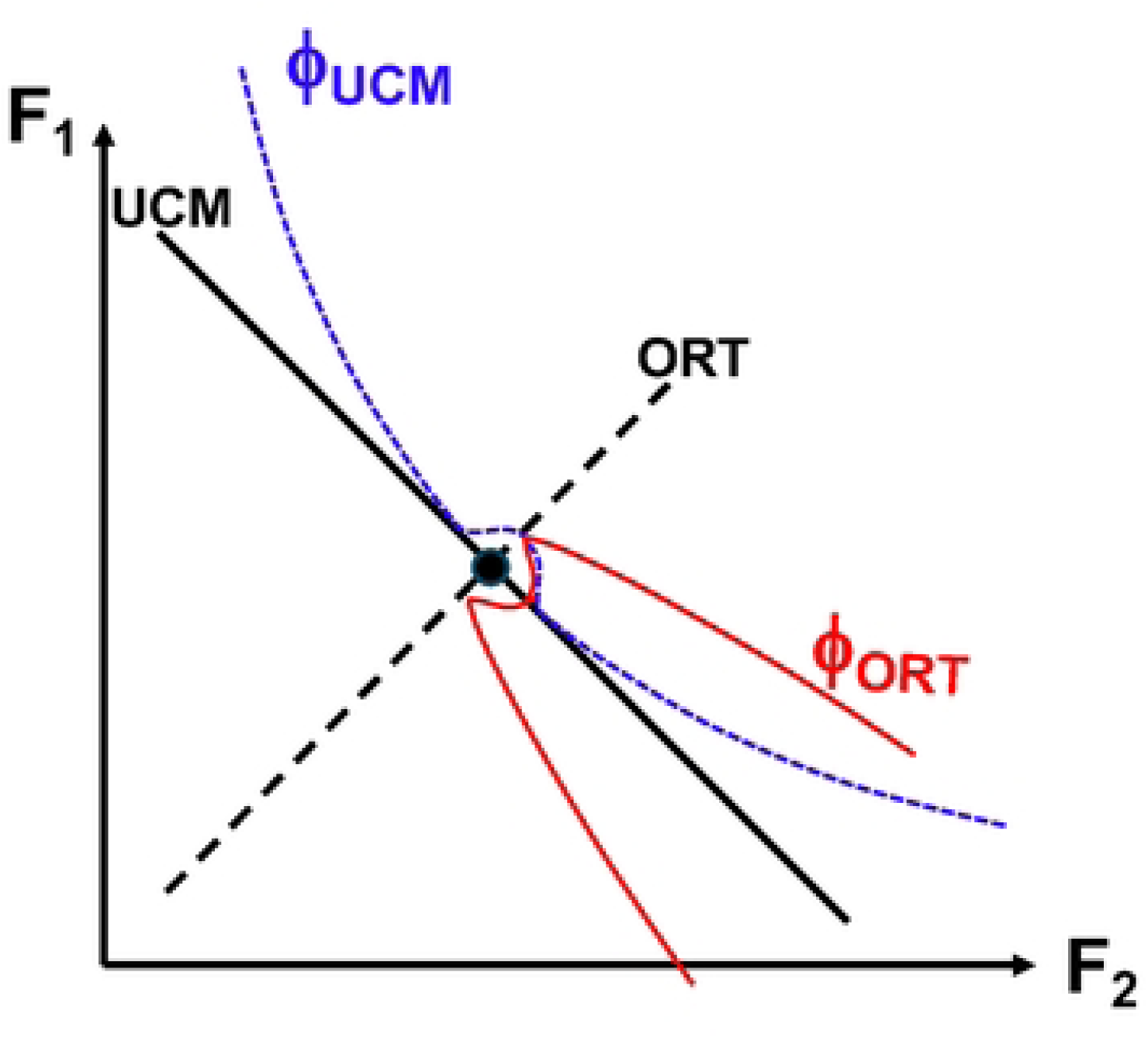
A schematic representation of a two-effector total force (F) production task (cf. Fig. 1). The uncontrolled manifold (UCM) is shown with the slanted solid line, and the ORT space with the dashed line. Different stability properties along the UCM and ORT are shown schematically with two potential fields, along the UCM (ϕ_UCM_, blue, dashed) and along the ORT (ϕ_ORT_, red, solid). Note the non-monotonic shape of the potential field, destabilizing within a small interval and stabilizing for larger deviations from the state (black dot).

The strong correlations between the H_Short_ values along the UCM and ORT directions observed across the participants for each of the visual feedback conditions have two potentially important implications. First, they represent, to our knowledge, the first experimental confirmation of coupling between processes within the UCM and ORT – an assumption made earlier based on analysis of force drifts (Ambike et al. 2016). Second, they suggest that healthy persons differ in characteristics of explorations within a relatively short range and these characteristics may represent a measure of one’s understanding the task and/or a personal trait (cf. De Freitas et al. 2019). Across populations, this index can turn into a useful clinical biomarker.

### Methodological issues and future developments

This is the first study exploring RW and drifts along both UCM and ORT. As in any first study, we had to make choices with respect to selecting specific parameters of the task and certain steps in the analysis. The current trial duration (60 s) was selected based on the first study that quantified RW and drifts along the UCM only (De et al. 2025a). An important question is whether characteristics of RW and drifts in individual participants are robust across repetitive trials. Given the relatively large number of conditions (three initial sharing patterns and four visual feedback conditions), to avoid fatigue, we have decided not to address this issue in this study and leave it for future exploration. The selection of specific sharing patterns, finger groups, and time windows for the analysis of the diffusion plots was motivated primarily by keeping consistency with the previous study. A number of issues remain unexplored including possible effects of hand dominance (cf. Sainburg 2005), symmetry of finger groups involved, processes within each pair of fingers, etc.

A non-trivial aspect of using sharing patterns different from 50:50 is the fact that the UCM for keeping the sharing index (SI) constant is not orthogonal to the UCM for keeping F_TOT_ constant. This implies that drifts along the UCM_SI_ have components along both ORT_F_ and UCM_F_ (the subscripts refer to the variable with respect to which the UCM is computed) potentially coupling the processes along these two spaces (cf. Ambike et al. 2016). Exploring a range of sharing patterns could shed light on the importance of this coupling for the observed drifts along the two subspaces defined for F_TOT_.

The persistence of RW at short time scales is compatible with the hypothesis that these processes can be defined primarily by the action of spinal circuitry including reflex feedback loops (as suggested in De et al. 2025a,b). This is corroborated by the finding that the instruction and visual feedback had only relatively minor effects on the values of H_Short_. In contrast, H_Long_ values were highly sensitive to visual feedback suggesting an important role of supraspinal structures in defining the stable (anti-persistent) portion of the potential field illustrated in Fig. 11. Note that the different roles of the spinal and supraspinal circuitry in performance-stabilizing synergies have been suggested based on studies of multi-effector (in particular, multi-finger) and intra-muscle synergies (Latash et al. 2023; De et al. 2024). Links between synergies and RW represent a potentially highly attractive and impactful topic for future analysis.

On a related topic, a recent study of patients with essential tremor performing accurate cyclical force production tasks has documented selective impairment of intra-muscle synergies (as compared to age-matched controls and patients with Parkinson’s disease) in contrast to unchanged multi-finger synergies (De et al. 2026). These observations have been interpreted as pointing at subtle changes at the level of spinal circuitry specific for essential tremor. The increased finger/hand tremor in patients with essential tremor suggests that this condition may be associated with a larger range of persistent RW resulting in larger oscillations between the borders separating the persistent and anti-persistent ranges (see Fig. 11), which could lead to larger-amplitude oscillations, i.e., tremor.

The frequency range of RW (mostly, between 10 and 25 Hz) suggests possible relations to physiological tremor (e.g., Vaillancourt and Newell 2000). To explore this relation, studies are needed involving a wide range of tasks and spaces of elemental variables. This can involve holding a joint configuration during a pointing task with and without visual feedback, as well as postural whole-body tasks similar to those explored earlier (Collins and De Luca 1993; Collins et al. 1995). Note that those earlier studies suggested much slower RW processes.

## References

Abolins V, Bernans E, Latash ML (2025) On the origin of variance along the uncontrolled manifold: Its effects on anticipatory and spontaneous changes in performance-stabilizing synergies in force production tasks. Neuroscience 569: 92–102.

Ambike S, Zatsiorsky VM, Latash ML (2015) Processes underlying unintentional finger force changes in the absence of visual feedback. Experimental Brain Research 233: 711–721.

Ambike S, Mattos D, Zatsiorsky VM, Latash ML (2016) The nature of constant and cyclic force production: Unintentional force-drift characteristics. Experimental Brain Research 234: 197–208.

Bernstein NA (1947) On the construction of movements. Medgiz: Moscow (in Russian). English translation is in: Latash ML (Ed.) (2020) Bernstein’s Construction of Movements. Routledge: Abingdon, UK.

Collins JJ, De Luca CJ (1993) Open-loop and closed-loop control of posture: a random-walk analysis of center-of-pressure trajectories. Experimental Brain Research 95: 308–318.

Collins JJ, De Luca CJ, Burrows A, Lipsitz LA (1995) Age-related changes in open-loop and closed-loop postural control mechanisms. Experimental Brain Research 104: 480–492.

Cuadra C, Corey J, Latash ML (2021) Distortions of the efferent copy during force perception: A study of force drifts and effects of muscle vibration. Neuroscience 457: 139–154.

De SD, Guysinsky M, Latash ML (2025a) Random walk and drifts within the uncontrolled manifold during multi-finger force production tasks. Discover Neuroscience 20: 28.

De SD, Hu X, Khan MS, Latash ML (2025b) Stability of performance in a hierarchical system during isometric force production and effects of visual feedback. Experimental Brain Research 244: 7.

De SD, Prado-Rico JM, Ricotta JM, De Jesus S, Seemiller J, Huang X, Latash ML (2026) Functional changes in spinal circuitry in essential tremor revealed with analysis of intra-muscle synergies. Journal of Neurophysiology (in press).

De SD, Ricotta JM, Benamati A, Latash ML (2024) Two classes of action-stabilizing synergies reflecting spinal and supraspinal circuitry. Journal of Neurophysiology 131: 152–165.

De Freitas PB, Freitas SMSF, Lewis MM, Huang X, Latash ML (2019) Individual preferences in motor coordination seen across the two hands: Relations to movement stability and optimality. Experimental Brain Research 237: 1–13.

Gelfand IM, Latash ML (1998) On the problem of adequate language in movement science. Motor Control 2: 306–313.

Goodman SR, Latash ML (2006) Feedforward control of a redundant motor system. Biological Cybernetics 95: 271–280.

Hausdorff JM, Peng CK, Ladin Z, Wei JY, Goldberger AL (1995) Is walking a random walk? Evidence for long-range correlations in stride interval of human gait. J Appl Physiol 78: 349–358.

Hultborn H, Brownstone RB, Toth TI, Gossard JP (2004) Key mechanisms for setting the input-output gain across the motoneuron pool. Progress in Brain Research 143:77–95.

Kitazawa S (2002) Optimization of goal-directed movements in the cerebellum: a random walk hypothesis. Neuroscience Research 43: 289–294.

Latash ML (2012) The bliss (not the problem) of motor abundance (not redundancy). Experimental Brain Research 217: 1–5.

Latash ML (2021) One more time about motor (and non-motor) synergies. Experimental Brain Research 239: 2951–2967.

Latash ML (2024) *Terra incognita* of the uncontrolled manifold. Journal of Neurophysiology 132: 1729–1743.

Latash ML, Madarshahian S, Ricotta J (2023) Intra-muscle synergies: Their place in the neural control hierarchy. Motor Control 27: 402–441.

Latash ML, Scholz JF, Danion F, Schöner G (2001) Structure of motor variability in marginally redundant multi-finger force production tasks. Experimental Brain Research 141: 153–165.

Latash ML, Scholz JP, Schöner G (2007) Toward a new theory of motor synergies. Motor Control 11: 276–308.

Latash ML, Shim JK, Smilga AV, Zatsiorsky V (2005) A central back-coupling hypothesis on the organization of motor synergies: a physical metaphor and a neural model. Biological Cybernetics 92: 186–191

Li S, Danion F, Latash ML, Li Z-M, Zatsiorsky VM (2001) Bilateral deficit and symmetry in finger force production during two-hand multi-finger tasks. Experimental Brain Research 141: 530–540

Mandelbrot BB, van Ness JW (1968) Fractional Brownian motions, fractional noises and applications. SIAM Rev 10: 422–437.

Martin V, Scholz JP, Schöner G (2009) Redundancy, self-motion, and motor control. Neural Computations 21:1371–1414

Martin V, Reimann H, Schöner G (2019) A process account of the uncontrolled manifold structure of joint space variance in pointing movements. Biological Cybernetics 113: 293–307.

Mitchell SL, Collins JJ, De Luca CJ, Burrows A, Lipsitz LA (1995) Open-loop and closed-loop postural control mechanisms in Parkinson’s disease: increased mediolateral activity during quiet standing. Neurosci Lett 197: 133–136

Parsa B, O’Shea DJ, Zatsiorsky VM, Latash ML (2016) On the nature of unintentional action: A study of force/moment drifts during multi-finger tasks. Journal of Neurophysiology 116: 698–708.

Parsa B, Terekhov AV, Zatsiorsky VM, Latash ML (2017) Optimality and stability of intentional and unintentional actions: I. Origins of drifts in performance. Experimental Brain Research 235: 481–496.

Rasouli O, Solnik S, Furmanek MP, Piscitelli D, Falaki A, Latash ML (2017) Unintentional drifts during quiet stance and voluntary body sway. Experimental Brain Research 235: 2301–2316.

Reschechtko S, Latash ML (2017) Stability of hand force production: I. Hand level control variables and multi-finger synergies. Journal of Neurophysiology 118: 3152–3164.

Reschechtko S, Zatsiorsky VM, Latash ML (2014) Stability of multi-finger action in different spaces. Journal of Neurophysiology 112: 3209–3218.

Roth AM, Calalo JA, Lokesh R, Sullivan SR, Grill S, Jeka JJ, van der Kooij K, Carter MJ, Cashaback JGA (2023) Reinforcement-based processes actively regulate motor exploration along redundant solution manifolds. Proceed Biol Sci 290: 20231475.

Sainburg RL (2005) Handedness: differential specializations for control of trajectory and position. Exercise and Sport Science Reviews 33: 206–213.

Scholz JP, Danion F, Latash ML, Schöner G (2002) Understanding finger coordination through analysis of the structure of force variability. Biological Cybernetics 86: 29–39.

Scholz JP, Schöner G (1999). The uncontrolled manifold concept: Identifying control variables for a functional task. Experimental Brain Research 126, 289–306.

Schwetlick L, Reich S, Engbert R (2025) Bayesian dynamical modeling of fixational eye movements. Biological Cybernetics 119: 13.

Vaillancourt DE, Newell KM (2000) Amplitude changes in the 8-12, 20-25, and 40 Hz oscillations in finger tremor. Clinical Neurophysiology 111: 1792–1801.

Vaillancourt DE, Russell DM (2002) Temporal capacity of short-term visuomotor memory in continuous force production. Experimental Brain Research 145:275–285

Vaz DV, Pinto VA, Junior RRS, Mattos DJS, Mitra S (2019) Coordination in adults with neurological impairment - A systematic review of uncontrolled manifold studies. Gait and Posture 69: 66–78.

Wilhelm L, Zatsiorsky VM, Latash ML (2013) Equifinality and its violations in a redundant system: Multi-finger accurate force production. Journal of Neurophysiology 110: 1965–1973.

Zhou T, Solnik S, Wu Y-H, Latash ML (2014) Unintentional movements produced by back-coupling between the actual and referent body configurations: Violations of equifinality in multi-joint positional tasks. Experimental Brain Research 232: 3847–3859.

Zhou T, Zhang L, Latash ML (2015) Intentional and unintentional multi-joint movements: their nature and structure of variance. Neuroscience 289: 181–193.

